# The Warburg Effect is the result of faster ATP production by glycolysis than respiration

**DOI:** 10.1101/2022.12.28.522160

**Authors:** Matthew A. Kukurugya, Denis V. Titov

## Abstract

Many prokaryotic and eukaryotic cells metabolize glucose to organism-specific byproducts instead of fully oxidizing it to carbon dioxide and water–a phenomenon referred to as the Warburg Effect. The benefit to a cell has been unclear, given that partial metabolism of glucose yields an order of magnitude less ATP per molecule of glucose than complete oxidation. We show that glycolysis produces ATP faster per gram of pathway protein than respiration in *E. coli*, *S. cerevisiae*, and mammalian cells. A simple mathematical model that uses yield, rate, and proteome occupancy of glycolysis and respiration as the only parameters accurately predicts absolute rates of glycolysis and respiration in all three organisms under diverse conditions. Our study suggests that the Warburg Effect is a consequence of the optimization of the rate of energy generation under the constraint of finite proteome space.

**One-Sentence Summary:** The Warburg Effect is a manifestation by which cells across kingdoms of life optimize the rate of energy production.

## Main Text

The Warburg Effect remains one of the most well-documented, yet incompletely understood phenomena in metabolism and cancer biology. In 1924, Otto Warburg made the seminal observation that *in vivo* tumors showed a preference for converting glucose to lactic acid via fermentation in the presence of oxygen instead of complete oxidation to carbon dioxide and water (*1, 2*). Although the initial discovery was made in tumor cells, the Warburg Effect-like metabolism has since been described for numerous proliferating cells, including acetate production through the Pta-AckA pathway in *E. coli,* ethanol fermentation in *S. cerevisiae,* and lactic acid fermentation in non-transformed mammalian cells. This phenomenon is also referred to as the Crabtree Effect in *S. cerevisiae* and “overflow metabolism” in *E. coli*. In addition, fast-twitch muscle cells–used for short and powerful bursts of activity–use fermentation for ATP production instead of respiration. To simplify nomenclature, we will collectively refer to various organism-specific pathways for partial metabolism of glucose (i.e., fermentation in *E. coli*, *S. cerevisiae*, and mammalian cells and respiro-fermentative Pta-AckA acetate pathway in *E. coli*) as *glycolysis*, to organism-specific pathways for complete oxidation of glucose to carbon dioxide and water as *respiration*, and to the preference for incomplete oxidation of glucose in the presence of oxygen as *the Warburg Effect*. Respiration can yield more than an order of magnitude more ATP per molecule of glucose than glycolysis. The rationale for why cells prefer lower ATP-yielding glycolysis for ATP production in the presence of oxygen has remained elusive.

Currently, no hypothesis is widely accepted to explain the occurrence of the Warburg Effect (*3*–*6*). Otto Warburg himself proposed that high glycolysis rates in cancer cells were due to an impairment in respiration (*2*). However, numerous studies have since demonstrated a prominent role of respiration in ATP production during cell proliferation (*7, 8*). Others have proposed that the transition from respiration to glycolysis in proliferating cells could be a mechanism to limit reactive oxygen species production (*9, 10*), to satisfy the demand for biosynthetic precursors (*11, 12*), to regulate the NADH:NAD^+^ redox balance (*13–15*), or to help yeast out-compete other microorganisms by rapidly consuming glucose (*16*). Our hypothesis builds on two sets of observations. First, a trade-off between yield and rate for ATP-producing pathways has been explored in theoretical studies, highlighting the advantages of high rate over high yield ATP-producing pathways under some conditions (*17, 18*). Second, several studies proposed competition for limited resources as an explanation for the Warburg Effect. In *E. coli*, it has been suggested that competition for plasma membrane area drives glycolysis, where glycolysis can produce more ATP per unit of limiting membrane area (*19*) since glucose transporters occupy less membrane space than the respiratory chain enzymes. Several studies proposed that glycolysis might be more proteome efficient than respiration, and a switch to glycolysis allows rapidly growing cells to allocate more of their proteome to ribosomes or other enzymes required for rapid growth (*20, 21*). Similarly, it has also been suggested that the diversity of glycolytic pathways in prokaryotes may represent a tradeoff between ATP yield and the enzyme mass needed to support the pathway flux (*22*).

The current hypotheses for why the Warburg Effect occurs have three limitations that we sought to address in this study. First, the current hypotheses assume that the Warburg Effect only occurs in proliferating cells, while it is well known that it can occur irrespective of cell growth (*23–26*). Second, most of the hypotheses are specific to one organism, which limits their ability to provide a unifying explanation for its occurrence in *E. coli*, *S. cerevisiae*, and mammalian cells. Finally, none of the proposed hypotheses can quantitatively predict absolute rates of glycolysis and respiration for a diverse set of organisms and conditions as would be required for a working theory.

Here we propose and test a unifying hypothesis for the occurrence of the Warburg Effect-like metabolism in *E. coli*, *S. cerevisiae*, and mammalian cells. We propose that the Warburg Effect is the expected result of the optimization of energy metabolism to allow maximal ATP production rate at different levels of glucose availability. To test our hypothesis, we developed a mathematical model that contains only five parameters, including yield and specific activity of respiration and glycolysis, and maximal proteome fraction that can be occupied by ATP-producing enzymes. We estimated the values of these five parameters from experiments (i.e., our model has no free parameters) and showed that our model quantitatively predicts the onset of the Warburg Effect, as well as glycolytic and respiration rates under various conditions in *E. coli*, *S. cerevisiae*, and mammalian cells irrespective of growth rate.

### Model that maximizes ATP production rate leads to the Warburg Effect

We propose a hypothesis that the Warburg Effect results from cells switching between glycolysis and respiration to achieve maximal ATP production rates at different environmental conditions. Cells have to obey three biochemical constraints while optimizing ATP production (Fig. 1A): i) glucose and oxygen consumption are limited by glucose and oxygen availability in the surrounding media, ii) a finite fraction of proteome is dedicated to ATP-producing enzymes while intracellular protein concentration remains constant as has been experimentally observed (*27–31*), and iii) the maximal ATP production rate by glycolysis or respiration pathways is limited by respective enzyme kinetics. To describe the maximal ATP production rate by respiration or glycolysis, we introduce a new metric that we call the specific activity of the pathway. We define the specific activity of the pathway as µmol of product produced (or substrate consumed) per min by mg of pathway protein by analogy with a widely used specific activity of enzymes. We assume that cells are free to shift the relative rates of glycolysis and respiration through a combination of changes in the expression of enzymes and transporters, post-translational modifications, and allosteric regulation as long as the above constraints are satisfied.

**Fig. 1.**
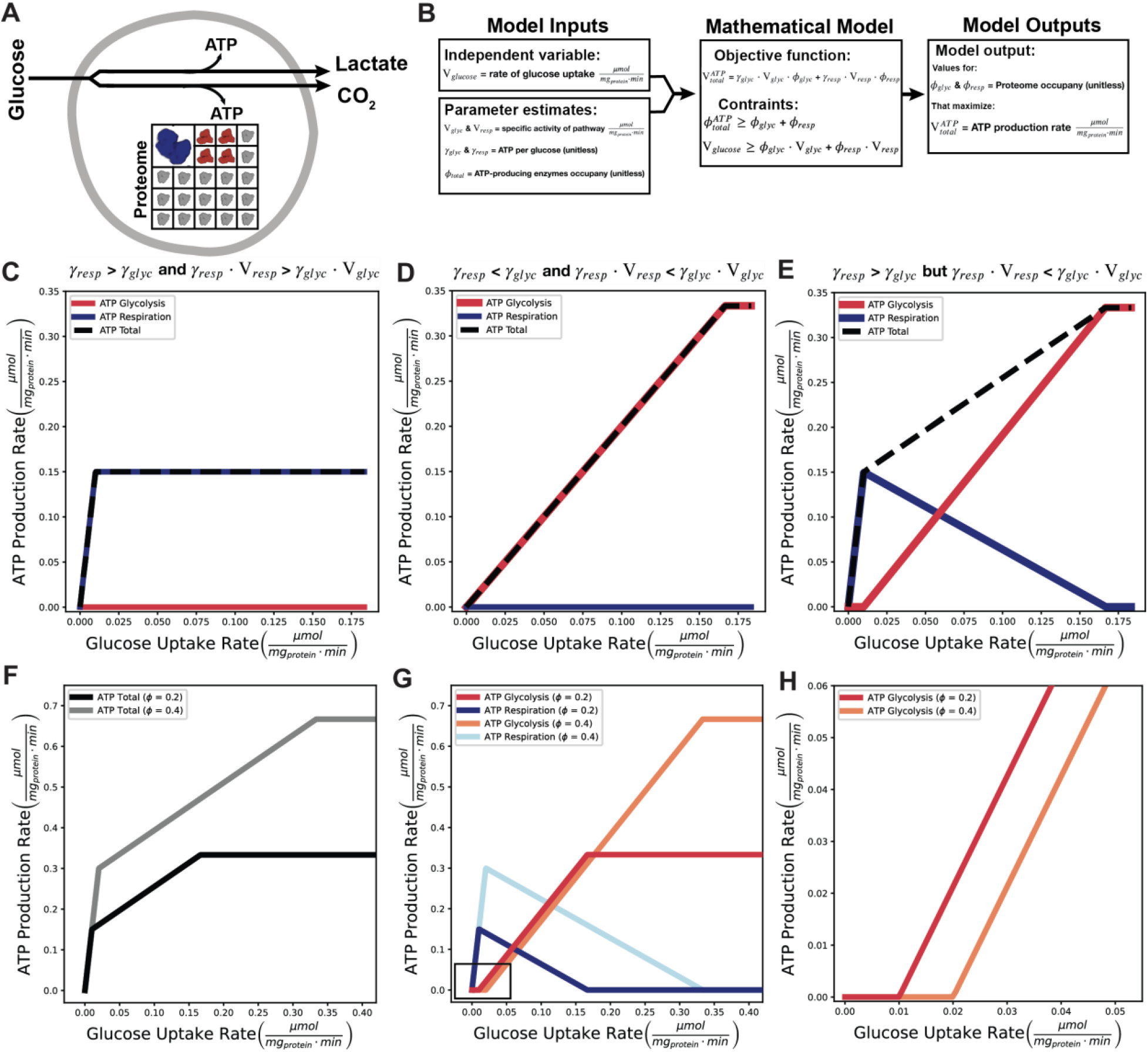
Model that maximizes ATP production rate leads to the Warburg Effect. **(A)** Illustration of model. **(B)** Overview of mathematical model. **(C)** Preferred ATP-producing pathways if parameters of yield of ATP per molecule of glucose (γ) and the specific activity of ATP production (*V* · *γ*) are both higher for respiration, **(D)** both higher for glycolysis, or **(E)** if the yield is higher for respiration (*γ*_*resp*_ > *γ*_*glyc*_), but the specific activity of ATP production is higher for glycolysis (*γ*_*resp*_ · *V*_*resp*_ < *γ*_*glyc*_ · *V*_*glyc*_). In each case, the total ATP production rate is represented by the dashed line. **(F)** Increasing the proteome space dedicated to ATP-producing enzymes increases ATP production rate across glucose uptake rates and **(G, H)** delays the switch from respiration to glycolysis. The box in Fig. 1G is shown in (H) to highlight the delayed onset of glycolysis.

To test the feasibility of our hypothesis, we constructed a mathematical model (Fig. 1B). Our aim was to keep the mathematical model as simple as possible to keep the interpretation of results straightforward. Our model uses linear programming to identify the ratio of glucose utilization through glycolysis and respiration that maximizes the ATP production rate of the cell subject to the constraints described in the last paragraph (see Fig. 1B for definitions of parameters):

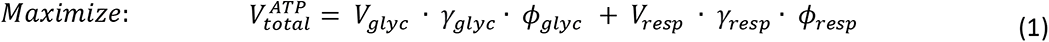

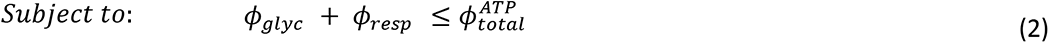

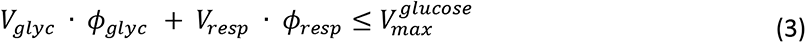

Our model predicts that if both the yield of ATP per molecule of glucose (*γ*) and the specific activity of ATP production (*V* · *γ*) are higher for respiration than glycolysis (e.i., *γ*_*resp*_ > *γ*_*glyc*_ and *V*_*resp*_ · *γ*_*resp*_ > *V*_*glyc*_ · *γ*_*glyc*_), then respiration will be the preferred ATP-producing pathway regardless of glucose availability as long as oxygen is available (Fig. 1, C and D). However, if the yield of ATP per molecule of glucose is higher for respiration (*γ*_*resp*_ > *γ*_*glyc*_), while the specific activity of ATP production is higher for glycolysis (*V*_*glyc*_ · *γ*_*glyc*_ > *V*_*resp*_ · *γ*_*resp*_), then respiration produces ATP faster at low glucose uptake rates, while glycolysis produces ATP faster at high glucose uptake rates (Fig. 1E). In other words, the cell can produce ATP at the fastest rate using high yielding respiration at low glucose availability (Fig. 1E and fig. S1, A to E), a mixture of high yielding respiration and high rate glycolysis at intermediate glucose availability (Fig. 1E and fig. S1B, C, F, G), or even with glycolysis alone to achieve the maximal ATP production rate (Fig. 1E and fig. S1, D and H).

While the specific values of the yield and specific activity predict the benefit of switching from respiration to glycolysis, the glucose uptake rate when the switch occurs is determined by the total proteome space dedicated to ATP-producing enzymes 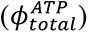 (Eq. 2). Without the proteome constraint, cells would always be able to express more respiratory proteins to increase ATP production without compromising on yield. Increasing the proteome fraction dedicated to ATP- producing enzymes 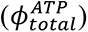 increases the capacity for ATP production across glucose uptake rates (Fig. 1F) and delays the switch from respiration to glycolysis (Fig. 1, G to H).

Our simple model assumes that glucose is used as the substrate for both glycolysis and respiration; however, the conclusions stay the same if a respiratory substrate other than glucose is used for respiration, such as acetate, glycerol, fatty acids, or amino acids (fig. S2 and Supplementary Discussion 1). The latter can be intuitively understood by considering that the majority of respiratory ATP is produced by tricarboxylic acid (TCA) cycle and electron transport chain (ETC) enzymes regardless of the specific substrate, and thus, the specific activity of respiratory ATP production is similar for different substrates.

In summary, our model predicts a transition from respiration to glycolysis as glucose availability increases (i.e., the Warburg Effect) under certain combinations of yield and specific activity of glycolysis and respiration.

### Glycolysis produces ATP at a faster rate than respiration

We next estimated the yields and specific activities of ATP production of glycolysis and respiration to determine if their values fall into the range where our hypothesis predicts the occurrence of the Warburg Effect: *γ*_*resp*_ > *γ*_*glyc*_ and *V*_*resp*_ · *γ*_*resp*_ < *V*_*glyc*_ · *γ*_*glyc*_. To test the general applicability of our hypothesis, we chose to focus on three organisms, *E. coli*, *S. cerevisiae*, and mammalian cells, since they span multiple kingdoms of life, exhibit Warburg Effect-like metabolic switch, have unique bioenergetic pathways (fig. S3), and have been extensively studied.

We first compiled the ATP yields of each pathway per molecule of glucose (fig. S4, A to C, Table S1). We found that respiration yields 10-, 8-, and 12-fold more ATP per glucose than glycolysis in *E. coli*, *S. cerevisiae*, and mammalian cells, respectively (fig. S4, A to C). In *E. coli*, we considered a third pathway, the respiro-fermentative Pta-AckA pathway (*32–34*), that uses glycolysis and ETC but not TCA cycle enzymes to produce acetate and carbon dioxide. The respiration yields 2-fold more ATP per glucose than the Pta-AckA pathway.

We used experimental data to estimate the specific activities of ATP production for each pathway (*γ* · *V*). To that end, we compiled an extensive dataset of physiological measurements to estimate the maximal rate of glycolysis and respiration per mg of cellular protein and divided those values by the fraction of the proteome occupied by glycolysis and respiration, respectively, determined from proteomics data (fig. S4, D to I, Table S2, Table S3). Our estimates showed that glycolysis has 0.3- (2- for Pta-AckA pathway), 2.4- and 6.1-fold faster rates per mg of pathway protein than respiration for *E. coli*, *S. cerevisiae*, and mammalian cells, respectively (Fig. 2, A to C). To the best of our knowledge, these are the only estimates of the specific activities of ATP production for any organism. Previously, Basan et al. (*21*) estimated the ratio of specific activity of ATP production of Pta-AckA pathway versus respiration in *E. coli* (called ratio of proteome efficiencies *ε*_*f*_/*ε*_*r*_ in that study) to be 1.5, similar to the lower bound of 95% confidence interval of our estimate of 1.6-2.3 (Fig. 2A).

**Fig. 2.**
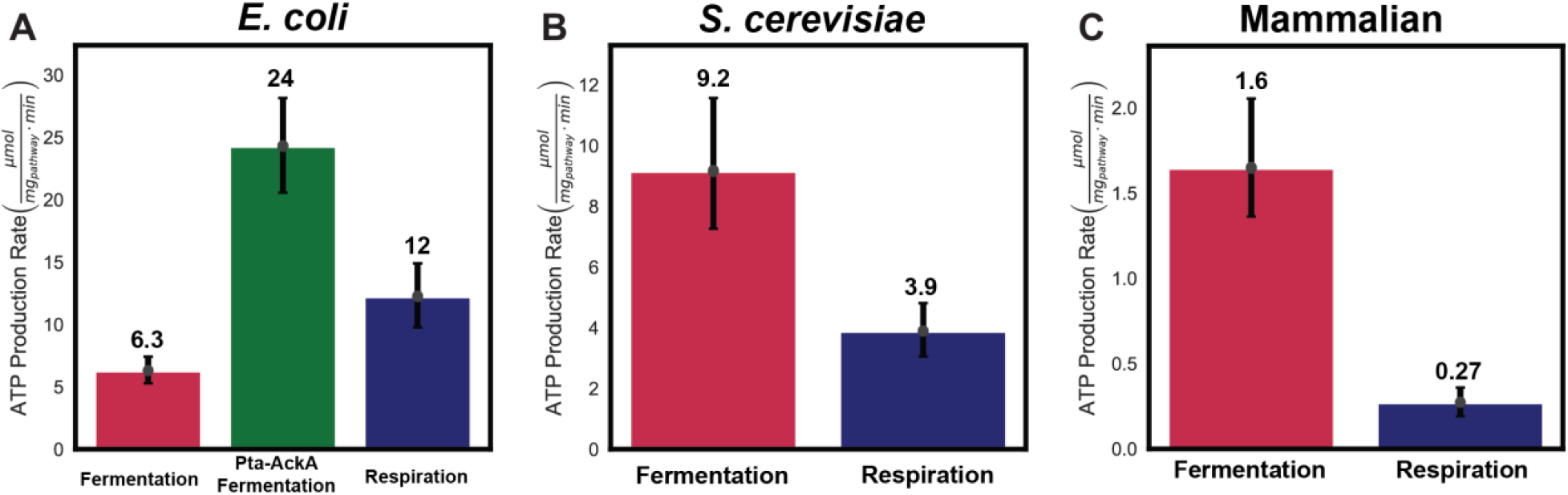
Pathway-specific ATP production rates for *E. coli*, *S. cerevisiae*, and mammalian cells. **(A)** Maximal ATP production rate (μmol mg_pathway_^-1^ min^-1^) for fermentative glycolysis (red), Pta-AckA pathway (green), and respiration (blue) for *E. coli*. (**B and C)** Maximal ATP production rate (μmol mg_pathway_^-1^ min^-1^) for fermentative glycolysis (red) and respiration (blue) for *S. cerevisiae* and mammalian cells, respectively. Error bars are the 95 percent confidence interval calculated from 10,000 bootstrap iterations.

In addition, we estimated the molecular weight of glycolysis and respiration by simply summing up the molecular weights of the individual components multiplied by their stoichiometry in the relevant pathway (fig. S4, J to L, Table S4). Respiration pathways were dramatically larger than glycolysis pathways in all three organisms (fig. S4, J to L, Table S4). Strikingly, the increase in the size of respiration in relation to glycolysis is larger than the increase in yield afforded by respiration for all pathways except for redox-neutral *E. coli* fermentation (fig. S4, M to O). These simple calculations corroborate our estimates from experimental data to suggest that the ATP production rate per mg of glycolysis proteins is larger than that of respiration.

Our estimates for ATP yield (*γ*_*resp*_ > *γ*_*glyc*_) and specific activity of ATP production (*V*_*resp*_ ·*γ*_*resp*_ < *V*_*glyc*_ · *γ*_*glyc*_) satisfied the parameter requirements in which our model predicts the switch from respiration to glycolysis at high glucose availability for respiro-fermentative Pta-AckA pathway in *E. coli*, and redox-neutral fermentative glycolysis in *S. cerevisiae* and mammalian cells. Importantly, our results do not predict a benefit for switching from respiration to fermentative glycolysis in *E. coli*. Thus, in addition to explaining why *E. coli*, *S. cerevisiae* and mammalian cells exhibit the Warburg Effect, our hypothesis provides an explanation for why *E. coli* exclusively use the respiro-fermentative Pta-AckA pathway in the presence of oxygen and not redox-neutral fermentative glycolysis as is the case for *S. cerevisiae*, and mammalian cells. Finally, we note that the specific activities of ATP production increase from mammalian cells to *E. coli* by over an order of magnitude for respiration and 4-5-fold for glycolysis, suggesting that there might be evolutionary pressure for microbes to increase the specific activity of ATP production.

### Model quantitatively predicts the onset of Warburg Effect

Having demonstrated that our hypothesis predicts the benefit of the Warburg Effect, we next tested whether our model can quantitatively predict the onset of the Warburg Effect as well as glycolysis and respiration rates measured under different experimental conditions in *E. coli*, *S. cerevisiae*, and mammalian cells. The additional model parameter that we needed for this test was the total fraction of the proteome dedicated to ATP-producing enzymes (*ϕ*_*total*_) that we estimated from proteomics data (fig. S4, D to F, Table S3). We used glucose uptake rate as the only input. For each given glucose uptake rate, the model used the four organism-specific model parameters as described in the last section (*γ*_*glyc*_, *γ*_*resp*_, *V*_*glyc*_, and *V*_*resp*_) and the total fraction of the proteome dedicated to ATP-producing enzymes (*ϕ*_*total*_) to determine the onset of the Warburg Effect as well as the glycolysis and respiration rates that allow cells to achieve a maximal ATP production rate (Fig. 3). We have compiled an extensive dataset of glucose uptake, glycolysis byproduct production, and respiration rates from 29 publications containing different strains, cell types, and experimental conditions to test the general applicability of our hypothesis (Table S5).

**Fig. 3.**
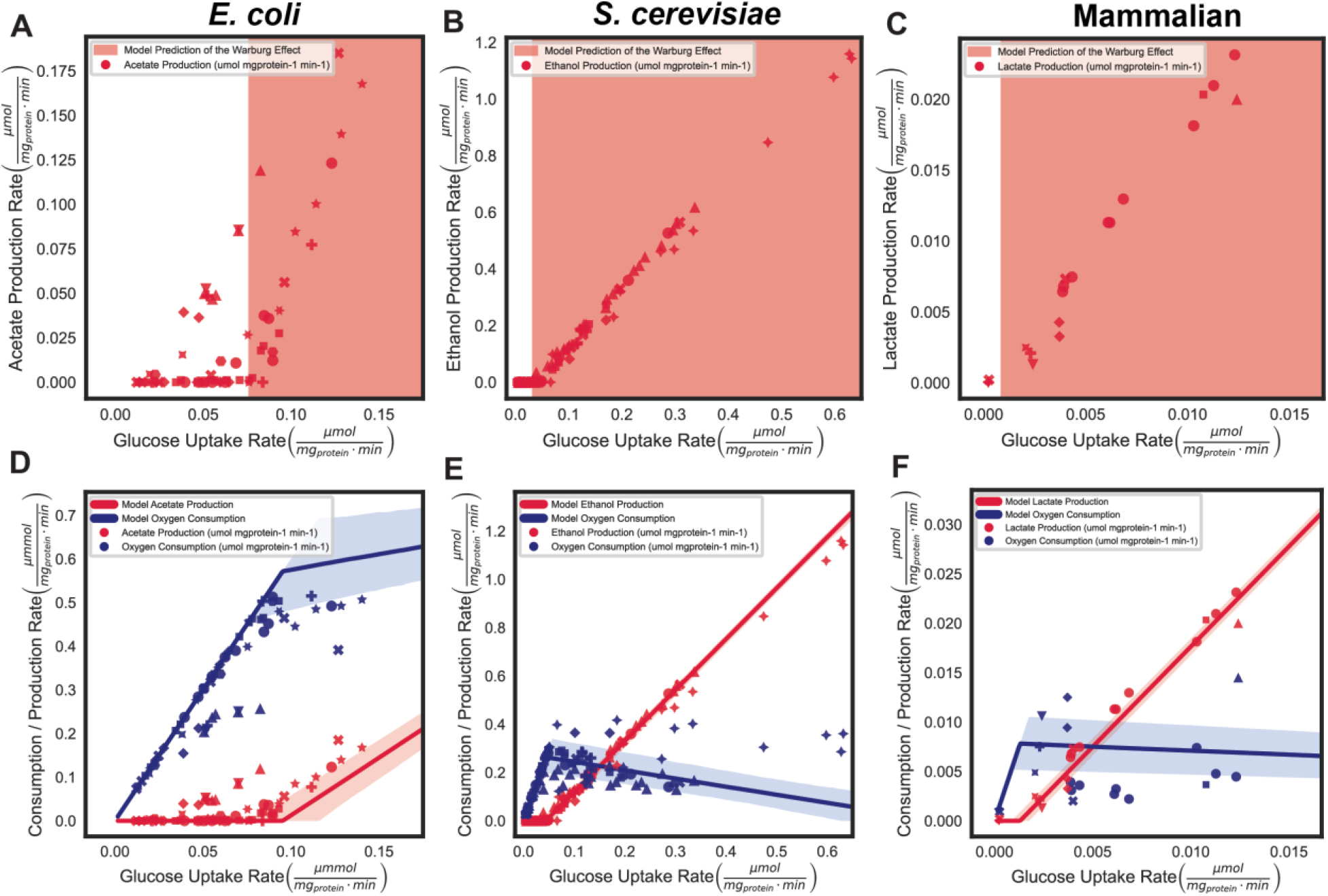
Organism-specific model predictions of the Warburg Effect. **(A,B,C)** Comparison of the model prediction (shaded region) for the glucose uptake rate in which glycolysis occurs **(D,E,F)** Comparison of model predictions (lines) and experimental observations (points) for glycolysis (red) and respiration (blue) rates of *E. coli*, *S. cerevisiae*, and mammalian cells, respectively. Note that each unique point shape represents data from a distinct publication.

Our model accurately predicts the glucose uptake rate range where the Warburg Effect is observed (Fig. 3, A to C). Furthermore, our model accurately predicted the absolute rates of acetate, ethanol, and lactate production, and oxygen consumption over two orders of magnitude of glucose consumption rates in *E. coli*, *S. cerevisiae,* and mammalian cells (Fig. 3, D to F). Importantly, the results in Figure 3 are not data fitting but predictions of the model using biochemical parameters estimated from independent experiments. Most of the experimental data are within the 95% confidence interval of model predictions, especially given the experimental errors of data points not displayed here for clarity. Our predictions for *S. cerevisiae* and mammalian cells were robust to different methods of accounting for mitochondrial proteome size in the estimation of *ϕ*_*total*_ and *V*_*resp*_ (fig. S5). Finally, we note that the change in total proteome allocation to ATP-producing enzymes would affect the ATP-producing capacity as well as the transition from respiration and glycolysis, as discussed above (Fig. 1, F to H). For our predictions, we used average ATP-producing space across many conditions for each organism, so the accuracy of our model predictions could be further improved by measuring ATP-producing space under relevant conditions, as it can change depending on cell type, nutrient availability, or disease state.

### Model predicts the onset of Warburg Effect independent of growth rate

Our hypothesis proposes that glucose availability and not the growth rate is driving the onset of the Warburg Effect. To tease apart the role of growth rate and glucose availability, we have used *E. coli* and *S. cerevisiae* datasets from nitrogen, and phosphorus-limited growth conditions where glucose uptake rate and growth rate diverge significantly (Table S5). In these datasets, the correlation between growth rate and glycolysis is weak (*ρ* = 0.50 and *ρ* = 0.41 for *E. coli* and *S. cerevisiae*, respectively) (Fig. 4, A and B) as compared to the correlation between the glucose uptake rate and glycolysis (*ρ* = 0.68 and *ρ* = 0.99, respectively) (Fig. 4, C and D). Our model accurately predicted the relationship between the glucose uptake and glycolysis rates in *S. cerevisiae* irrespective of growth rate (Fig. 4D). For *E. coli*, our model overestimated the shift to acetate production under nitrogen-limited conditions (yellow points) (*37*). We speculate this is due to the known preference of *E. coli* to decrease the proteome fraction allocated to ATP-producing enzymes *ϕ*_*total*_ at high glucose availability (*21, 36*). The latter invalidates the constant *ϕ*_*total*_ assumption of our simple model under these specific conditions in *E. coli,* but is entirely consistent with our overall hypothesis that *E. coli* switched to the Pta-AckA glycolysis pathway because of its higher specific activity of ATP production in relation to respiration. These results suggest that, under nitrogen- and phosphorus-limited conditions in *E. coli* and *S. cerevisiae*, glycolysis is utilized as a major ATP production strategy as compared to the glucose-limited culture with the same growth rate due to the excess availability of glucose (Fig. 4, C and D). A hypothesis dependent on growth rate instead of glucose availability would not capture this phenomenon.

**Fig. 4.**
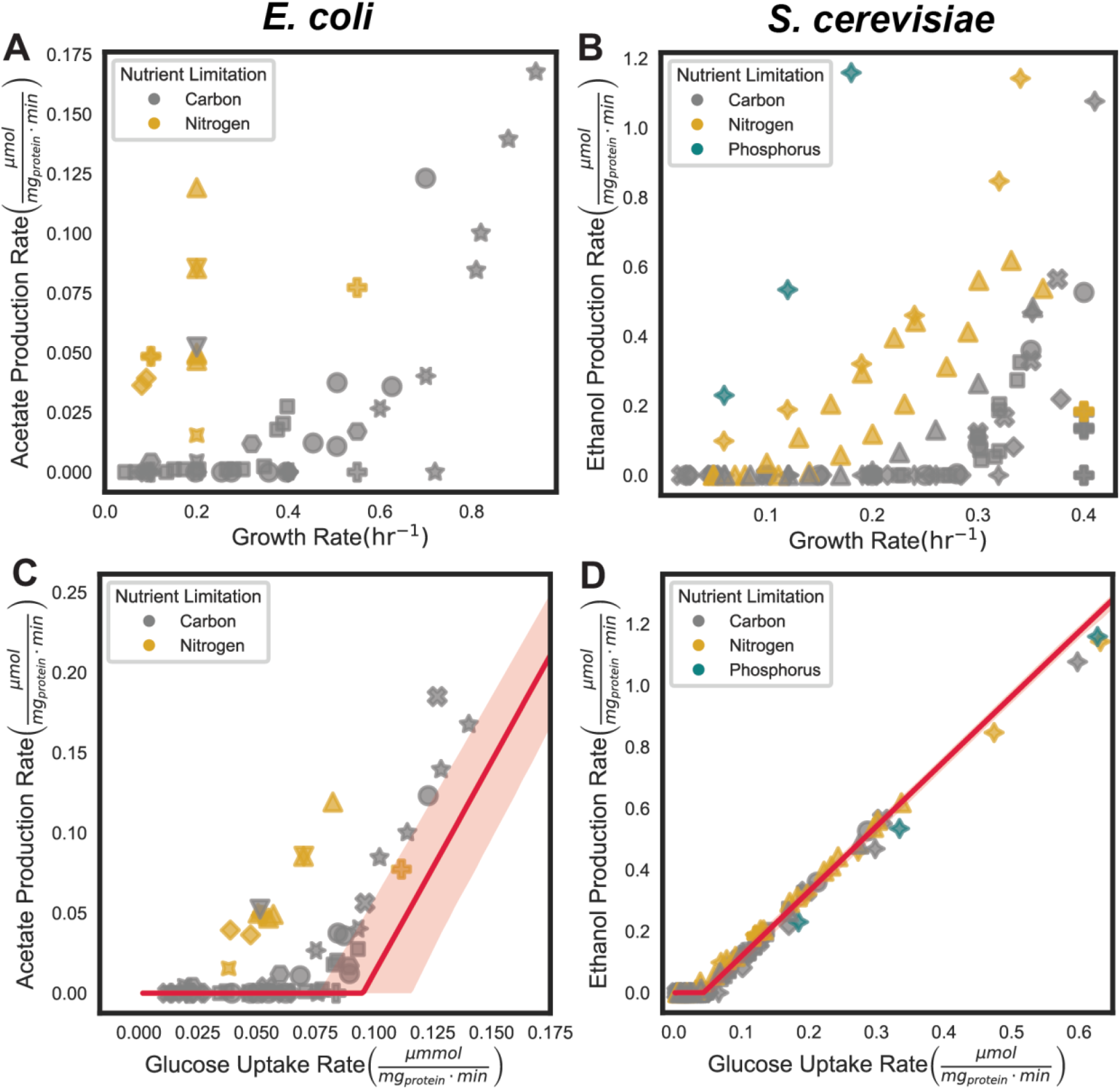
The Warburg Effect occurs independent of growth rate. **(A)** *E. coli* growth rate (hr^-1^) vs. the observed acetate production rate (μmol per mg cellular protein per min) or **(B)** glucose consumption rate (μmol per mg cellular protein per min) for carbon- and nitrogen-limited cultures (grey and yellow, respectively). **(C)** *S. cerevisiae* growth rate (hr^-1^) vs. the observed ethanol production rate (μmol per mg cellular protein per min) or **(D)** glucose consumption rate (μmol per mg cellular protein per min) for carbon-, nitrogen-, phosphorus-limited cultures (grey, yellow, green, respectively). Note that each unique point shape represents data from a distinct publication.

## Conclusion

In summary, we propose and test a hypothesis that the Warburg Effect arises from the optimization of energy metabolism in which the cell maximizes the ATP production rate for a given glucose availability. Our results suggest that while respiration has a higher yield of ATP per molecule of glucose, the ATP production rate per pathway mass is higher for Pta-AckA pathway in *E. coli*, and redox-neutral fermentative glycolysis in *S. cerevisiae*, and mammalian cells. In other words, glycolysis is more compact than respiration, which allows glycolysis to produce ATP faster than respiration when both pathways occupy the same fraction of the proteome and excess glucose is available. Our hypothesis also explains the preferential use of the Pta-AckA pathway in *E. coli* as only the Pta-AckA pathway, and not redox-neutral fermentative glycolysis, can produce ATP at a faster rate than respiration in *E. coli*. The ability of our model to make quantitative predictions using only five experimentally determined parameters provides a path for further validation of our hypothesis with additional experiments to improve estimates of the five model parameters and to generate more data connecting glucose uptake rate to glycolysis and respiration rates under different experimental conditions. Finally, we believe that our hypothesis can be used to explain a host of other observations beyond the Warburg Effect and Pta-AckA pathway preference, including the use of glycolysis by fast-twitch muscle cells (*25, 26*) and the use of proton-pumping (high yield) and non-proton pumping (high rate) respiratory chain components under different conditions that are ubiquitous among microbes.

## Acknowledgments

We thank A. Arkin, B. Bennett, D. Botstein, A. Flamholtz, H. Garcia, R. Heald, J. Thorner, R. Wang, X. Yang and members of the Titov Lab for valuable comments on the manuscript.

## Author contributions

Conceptualization: MAK, DVT

Methodology: MAK, DVT

Investigation: MAK, DVT

Data Curation: MAK, DVT

Software: MAK, DVT

Visualization: MAK, DVT

Funding acquisition: DVT

Writing – original draft: MAK, DVT

Writing – review & editing: MAK, DVT

## Data and materials availability

All data are available in the main text or the supplementary materials. All data and code used in the figure generation are available as a GitHub repository via https://github.com/DenisTitovLab/WarburgEffectModel

## Supplementary Materials

### Materials and Methods

#### Data curation

All data used in this report are from the existing primary literature. Details of the experimental conditions as well as exact steps taken to standardize the measurements can be found in the corresponding Supplementary Tables for each organism and measurement type.

#### Analysis code and figure generation

All data and code used in the figure generation are available as a GitHub repository via https://github.com/DenisTitovLab/WarburgEffectModel

#### Estimation of proteome occupancy by glycolysis and respiration

We utilized a total protein approach and previously published measurements of the absolute quantification of proteins using mass spectrometry to estimate the fraction of the proteome occupied by glycolysis ϕ_glyc_ and respiration ϕ_resp_ (Table S3). Only studies that used 8M urea or sodium dodecyl sulfate in the lysis buffer to improve extraction of membrane proteins were included. Total protein approach assumes that a protein’s abundance within a cell’s proteome is proportional to that protein’s mass spectrometry (MS) signal over the total MS signal (Eq. S1, Table S6).

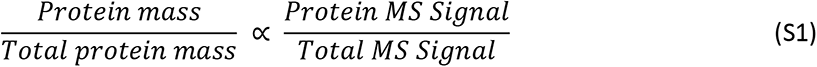

This approach was previously validated by showing that it can accurately quantify proteins within their expected physiological range spanning five orders of magnitude.

ETC complexes are imbedded in the mitochondrial membrane and, therefore, are often extracted with less efficiency. The level of ETC complex subunits with greater than 20 percent missing values were corrected using the levels of other subunits in the same complex on a per sample basis. Both subunit mass and stoichiometry were accounted for in the correction.

The specific enzymes included in the calculation of ϕ_glyc_ and ϕ_resp_ for *E. coli*, *S. cerevisiae*, and mammalian cells are included in Supplementary Table (Table S7). To match the respective physiological measurements for *E. coli* and *S. cerevisiae*, ϕ_glyc_ was only calculated for fermentable carbon substrates (i.e. sugars) and ϕ_resp_ is only calculated for non-fermentable carbon substrates (i.e. acetate, pyruvate, succinate, glycerol, fumarate) both in batch culture. For the respiro-fermentative Pta-AckA pathway in *E. coli*, ETC components were included to account for redox balance and additional ATP production.

For respiration in *S. cerevisiae* and mammalian cells, a list of core respiratory enzymes from the TCA cycle and ETC were included in the calculation of ϕ_resp_ (Table S7). In addition, any mitochondrial protein that positively correlated was correlated with the expression of the sum of the core respiration proteins (*ρ > 0)* and was statistically significant after Benjamini-Hochberg correction (p < 0.05). The statistical significance represents a two-tailed p-value, which is the probability of a correlation coefficient at least as big to be observed if the null hypothesis is true. All proteins included in the ϕ_resp_ for *S. cerevisiae* and mammalian cells are listed in Supplementary Table S8.

To account for the differences in protein expression in minimal and complete media for *E. coli* and *S. cerevisiae*, we subtracted the fraction of the pathways utilized for biosynthesis in minimal media. We estimated the fraction of glycolysis and the TCA cycle used for biosynthesis by calculating the average percent of glucose that becomes biomass in minimal media (Table S8). The fraction of the proteome occupied by glycolytic and TCA cycle enzymes was multiplied by (1 - biomass yield) to get the fraction of glycolytic and TCA cycle enzymes used for energy generation. For *E. coli*, the fraction of glycolytic and TCA cycle enzymes that was utilized for biomass production was 42% on fermentative substrates and 41% for respiratory substrates. For *E. coli*, minimal media accounted for 38 of the 55 utilized conditions. For *S. cerevisiae*, the fraction of glycolytic and TCA cycle enzymes that was utilized for biomass production was 14% on fermentative substrates and 53% for respiratory substrates. For *S. cerevisiae*, minimal media accounted for 14 of the 50 utilized conditions.

#### Estimation of maximal cellular activity of glycolysis and respiration

Absolute specific activities of glycolysis and respiration were estimated from experiments in which the maximal activity of each pathway was measured. For *E. coli, S. cerevisiae*, and mammalian cells, the maximal acetate, ethanol, and lactate production rates were used for glycolysis, respectively (Table S2). For *E. coli and S. cerevisiae*, these values were taken from measurements grown in batch culture on a fermentable substrate. For respiration, the maximal oxygen consumption rates were utilized for *E. coli, S. cerevisiae*, and human cells (Table S2). For *E. coli* and *S. cerevisiae*, these values were taken from measurements grown in batch culture or accelerostat on a nonfermentable substrate. Each value was converted to units of μmol per mg cellular protein per min. Note that for human cells, all oxygen consumption is from seahorse assays in which FCCP–- Antimycin A (or Rotenone), which represents the maximal respiratory capacity without non-mitochondrial respiration.

For studies that reported values in units consisting of gram cell dry weight (gCDW), it was assumed that 55% and 44% of that dry cell mass consisted of protein in *E. coli* and *S. cerevisiae,* respectively (Table S9). The estimates of gCDW per g protein are based on the studies for *E. coli* and *S. cerevisiae* compiled in Table S10. For mammalian cells studies which reported values in units of cell number, representing 9 of 14 studies, it was assumed that each cell contained 300 pg of protein (*1–3*).

#### Estimation of the molecular weight of glycolysis and respiration pathways

Molecular weights of the pathways were calculated by summing the molecular weights of all proteins in each of the pathways corrected for pathway stoichiometry (Table S4). For each enzyme that has isoforms, the average molecular weight was used. For each enzyme that is a multi-subunit protein complex, the sum of the subunits was used with subunit stoichiometry accounted.

#### Estimation of ATP yield per glucose of glycolysis and respiration

The ATP yield for (γ_resp_ and γ_glyc_) was estimated for *E. coli*, *S. cerevisiae*, and mammalian cells. Fermentative glycolysis yield of 2 ATP per glucose is well-defined for all three organisms (fig. S4, A to C and Table S1).

To define the theoretical yield of respiration for each organism, experimentally derived measurements (*4–6*) in combination with the following calculations were used to account for ATP produced from glycolysis, the TCA cycle, and the ETC (Table S1). In glycolysis, glucose is converted to pyruvate, producing a net gain of 2 ATP and 2 NADH per glucose. In *S. cerevisiae* and mammalian cells, the NADH produced from glycolysis are shuttled into the mitochondria to enter the ETC at a cost of 1 H^+^ per NADH. In the TCA cycle, pyruvate is further oxidized, generating a total of 8 NADH, 2 FADH_2_, and 2 ATP per glucose. In the ETC, NADH and FADH_2_ are oxidized to generate a proton gradient in the inner mitochondrial membrane (or plasma membrane in *E. coli*). The proton gradient is then used to drive the production of ATP synthase. Each organism has a unique number of H^+^ translocated per NADH and FADH_2_ based on the translocated H+ per electron ratio of the utilized ETC components. For *E. coli*, the ratio of translocated H^+^ per electron is 2 for NADH:ubiquinone oxidoreductase and 2 for cytochrome bo oxidase. For *S. cerevisiae*, the ratio of translocated H^+^ per electron is 0 for NADH dehydrogenase, 1 for cytochrome c reductase, and 2 for cytochrome c oxidase. For mammalian cells, the ratio of translocated H^+^ per electron is 2 NADH:ubiquinone oxidoreductase (CI), 1 for cytochrome c reductase (CIII), and 2 for cytochrome c oxidase (CIV). Note that each NADH or FADH_2_ donates two electrons. Therefore, the total number of translocated H^+^ is calculated by the following equation where mammalian cells are used as an example:

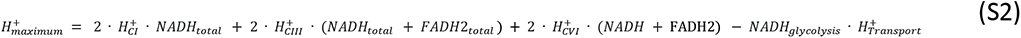

However, it is well-known that proton leak occurs in the membrane. For mammalian cells, the proton leak was calculated from changes in the oxygen consumption rate measured in which oligomycin–- antimycin A (or rotenone) was measured (Table S10). We determined that the proton leak was 25%. For *E. coli* and *S. cerevisiae*, we assumed ∼75% of theoretical maximal yield to account for proton leak. Furthermore, the ratio of H^+^/ATP is 4, 3, and 3 for *E. coli*, *S. cerevisiae*, and mammalian cells, respectively (*4–6*). In addition, the cost of transporting each ATP from the mitochondria is 1 H^+^. Therefore, the total number of ATP produced through respiration is calculated by:

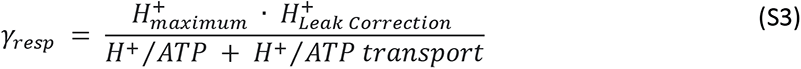

The ATP yield of respiration was determined to be 20, 16, and 24 ATP per molecule of glucose for *E. coli*, *S. cerevisiae*, and mammalian cells, respectively.

The Pta-AckA pathway in *E. coli* utilizes the ETC for oxidation of the 4 NADH produced per molecule of glucose. Therefore, the same approach was applied for this pathway.

#### Estimation of the specific activity of ATP production of glycolysis and respiration

We used experimental data to estimate the specific activities of ATP production of glycolysis and respiration (fig. S4, G to I, Table S2). The specific activity of glucose consumption was calculated by taking the maximal cellular glucose consumption rate by a given pathway (µmol glucose per mg of cellular protein per min) and dividing it by the fraction of the proteome occupied by the pathway (*ϕ*_*glyc*_ or *ϕ*_*resp*_).

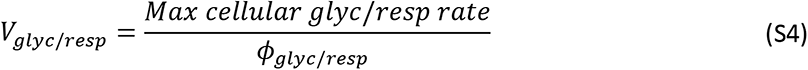

We determined the maximal glycolysis and respiration rates for *E. coli, S. cerevisiae*, and mammalian cells by compiling an extensive dataset from 38 independent publications containing 57 measurements of production rate of glycolysis products (i.e, acetate, ethanol, lactate) and oxygen consumption rates, which were converted to glucose consumption rate using known stoichiometry of the pathways (Table S2). For each organism, we calculated *ϕ*_*glyc*,_ and *ϕ*_*resp*_ from proteomics data (fig. S4, D to F; Table S3). As described in the *Estimation of proteome occupancy by glycolysis and respiration* section, we made corrections to *ϕ*_*resp*_ to account for mitochondrial proteins that are required for mitochondrial biogenesis and function but are not core respiration components by including into *ϕ*_*resp*_ mitochondrial proteins whose expression was significantly (p < 0.05) and positively correlated (*ρ > 0*) with the sum of the TCA and ETC proteins. In addition, we corrected proteomics data for *E. coli* and *S. cerevisiae* grown in minimal media by subtracting fractions of glycolysis and the TCA cycle that are used for biosynthesis. We note that this estimation procedure is conservative as membrane-bound ETC complexes are expected to be underrepresented in proteomics data. We also did not correct for the use of glycolysis and TCA cycle for biosynthesis in rich media, and we did not include all mitochondrial proteins in *ϕ*_*resp*_ . Correcting for any of these observations would further increase the ratio of specific activities of ATP production by glycolysis to respiration.

#### Mathematical Model

To describe the Warburg Effect, we use a linear programming optimization model. The objective function of the model is to maximize the ATP production rate given the constraints as described in (Eq. 1), (Eq. 2), and (Eq. 3) reproduced here for convenience:

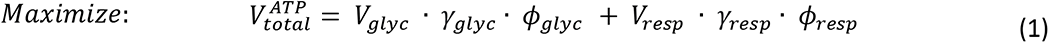

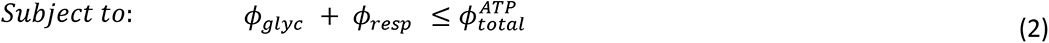

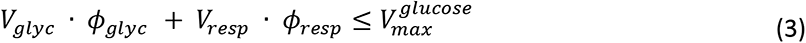

For each explicitly stated glucose uptake, the model assigns the values for ϕ_glyc_ and ϕ_resp_ which maximize the ATP production rate (Eq. 1) and satisfy the constraints (Eq. 2 and Eq. 3). The ϕ_glyc_ and ϕ_resp_ values represent the proteome fractions of glycolysis and respiration that are being expressed and utilized. Note that the expression of a given pathway can exceed its utilization, but never the inverse. Using the assigned ϕ_glyc_ and ϕ_resp_ values, the rate of acetate, ethanol, or lactate production (Eq. S5) and oxygen consumption (Eq. S6) are calculated for *E. coli*, *S. cerevisiae*, and mammalian cells:

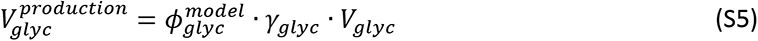

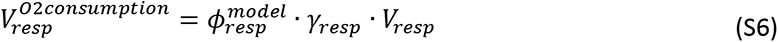

To estimate the confidence intervals of model prediction, we performed bootstrapping to resample the five parameter estimates (i.e., fraction of the proteome occupied by pathway, specific activity, and ATP yield) with replacement (N=10,000). After each round of sampling, the specific activity of each pathway and total proteome occupancy of ATP-production enzymes is calculated as described previously. The sampled parameter values for proteome occupied by pathway, specific activity, and ATP yield from each round of bootstrapping were then used in the linear program described above. All 10,000 results were stored in a data table. The mean and 95% confidence intervals were calculated and plotted. For *S. cerevisiae* and mammalian cells, two additional estimates were conducted: (1) with no additional mitochondrial proteins and (2) with all mitochondrial proteins (fig. S5, Table S8).

### Supplementary Text

#### Theoretical Model of Respiratory Substrate Utilization and Co-utilization with Glucose

We have extended our mathematical model to explore the effect of adding a respiratory substrate other than glucose.

The only differences between the core model and extended model are that we decompose the original *V_resp_*, *γ_resp_*, and *ϕ_resp_* into a contribution from glucose 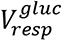, 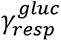, and 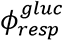 and a contribution from alternative respiratory substrate 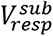, 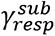, and 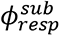, and we modify the equations and constraints of the core model to include these new terms and add a new constraint for the maximal respiratory substrate uptake:

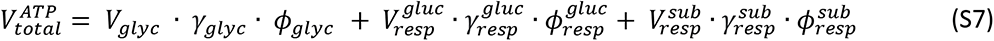

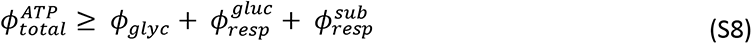

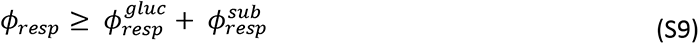

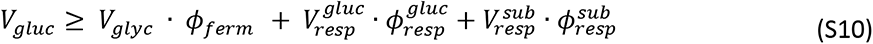

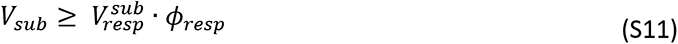

We assumed that the specific activity of respiratory substrate utilization in units of μmol per mg cellular protein per min are the same for both glucose and the respiratory substrate as this value should be mostly set by the shared pathways of TCA cycle and ETC. Our simulations show that even in the presence of saturating respiratory substrate, there is no effect of additional respiratory substrate on the ratio of glycolysis and respiration used by the cell at all glucose uptake rates except for the very low glucose uptake rate when respiration is not saturated by glucose (fig. 2, A to B). Together, our results suggest that the excess of glucose that can be diverted to glycolysis, not the presence of a respiratory substrate, drive the Warburg Effect.

**Fig. S1.**
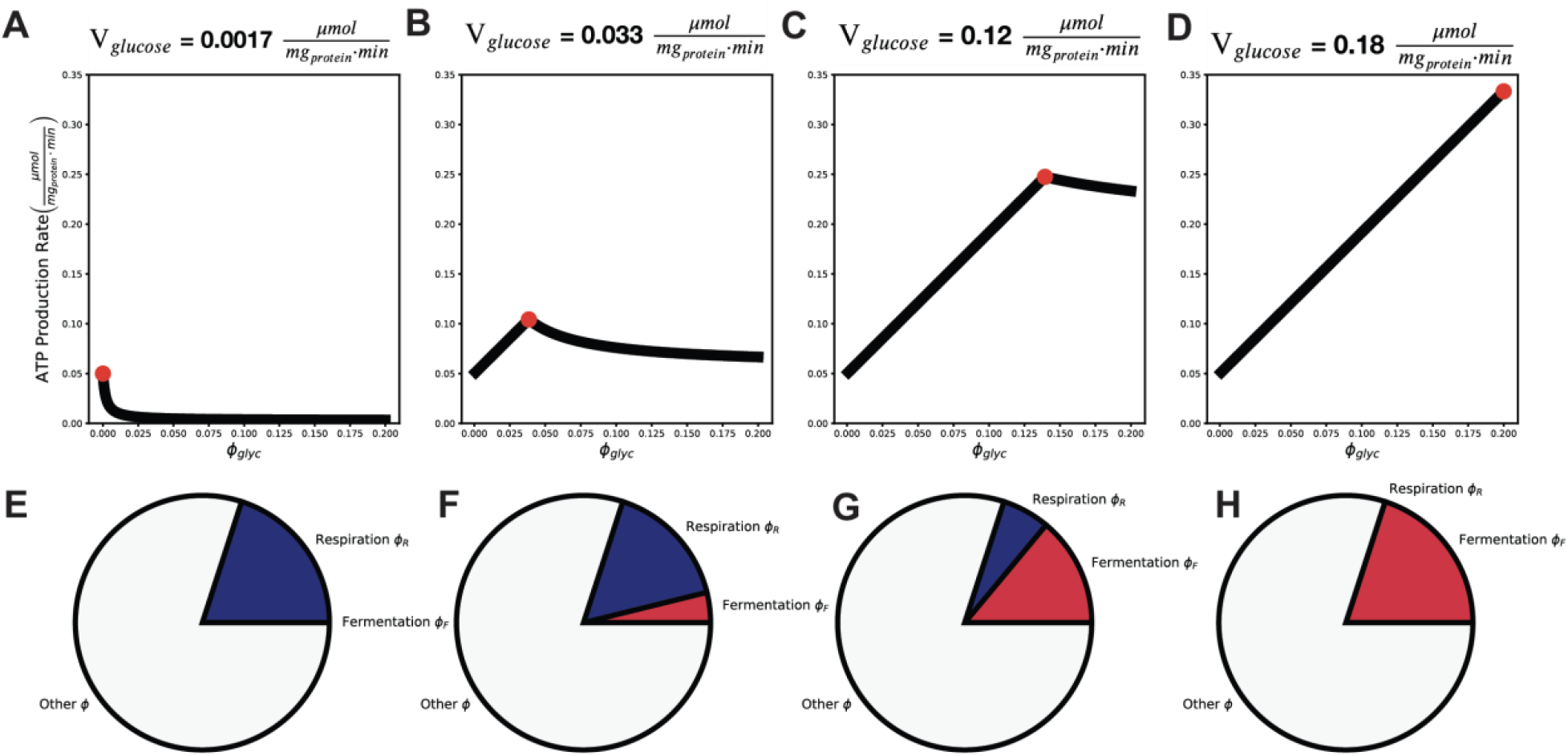
Model predictions under varying glucose uptake rates. **A)** Fraction of glycolysis that yields the highest ATP production rate (highlighted with a red dot) for a glucose uptake rate *γ*_*gluc*_ = 0.0017 μmol per mg cellular protein per min. **B)** *γ*_*gluc*_ = 0.033 μmol per mg cellular protein per min. **C)** *γ*_*gluc*_ = 0.12 μmol per mg cellular protein per min. **D)** *γ*_*gluc*_ = 0.18 μmol per mg cellular protein per min. **E, F, G, H)** fractions of glycolysis and respiration occupying the ATP producing space of the proteome under the same conditions as a), b), c) and d), respectively.

**Fig. S2.**
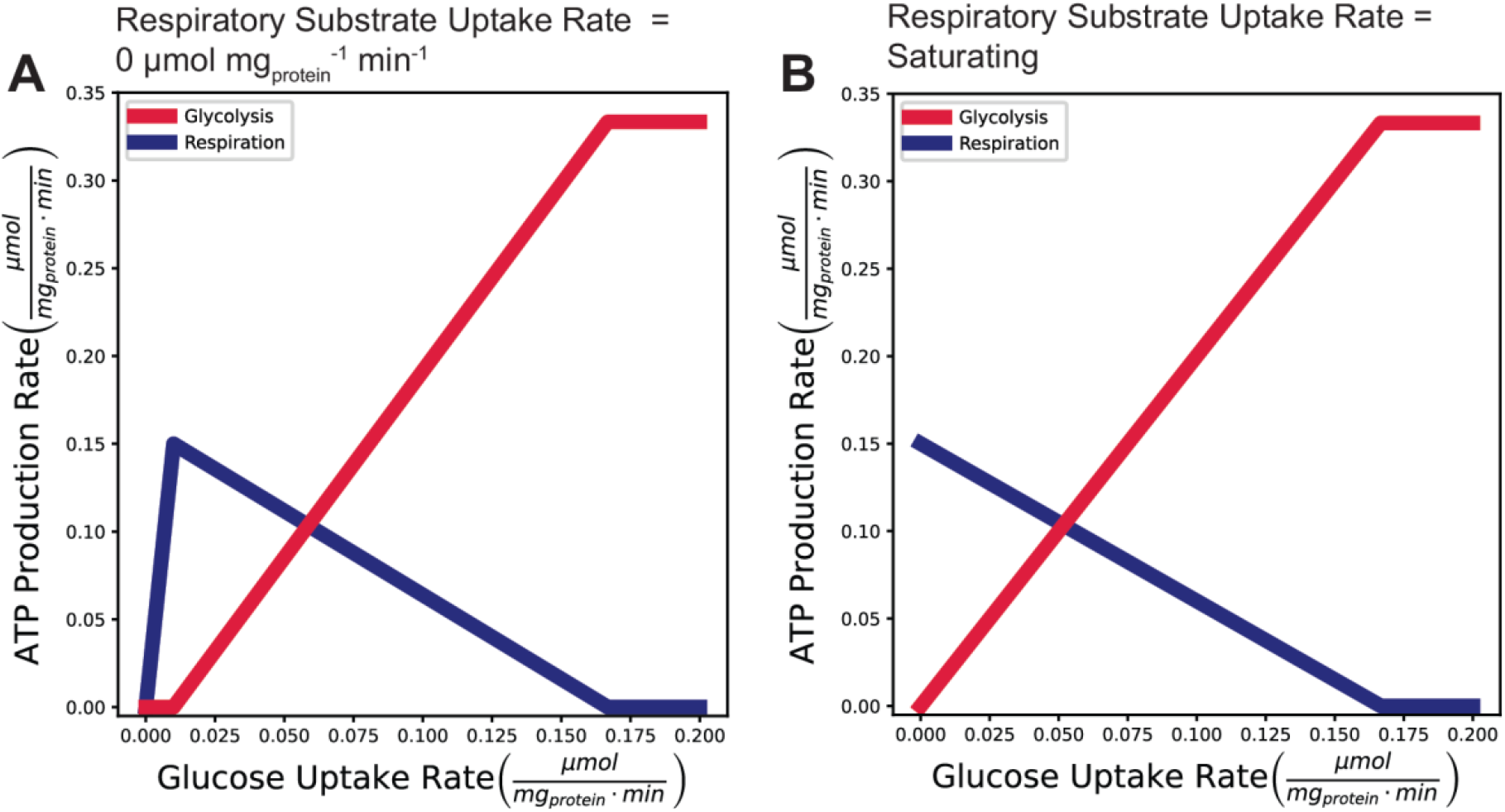
Model predictions under varying glucose and respiratory substrate uptake rates. **A)** ATP production rates of glycolysis (red) and respiration (blue) a) in the absence of respiratory substrate and **B)** in the presence of saturating respiratory substrate.

**Fig. S3.**
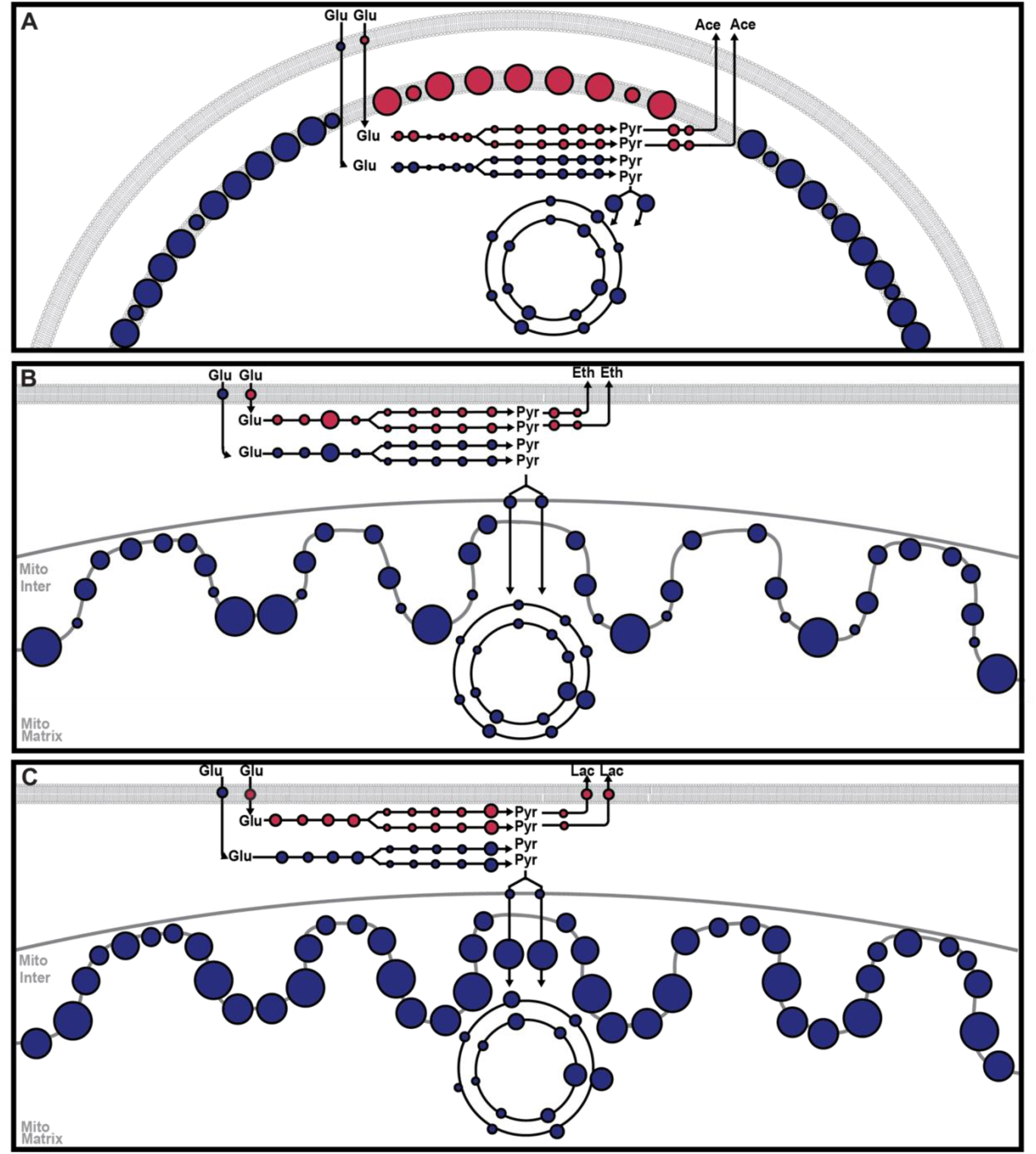
Glycolysis and respiration pathway sizes in E. coli, S. cerevisiae and mammals. **A-C)** Glycolysis (red) and respiration (blue) pathway outline for **A)** *E. coli* (respiro-fermentative Pta-AckA pathway that uses both glycolysis and ETC is displayed), **B)** *S. cerevisiae*, and **C)** mammalian cells. Size of circle is proportional to the MW of each enzyme in the pathway. Furthermore, the approximate pathway stoichiometry is represented.

**Fig. S4.**
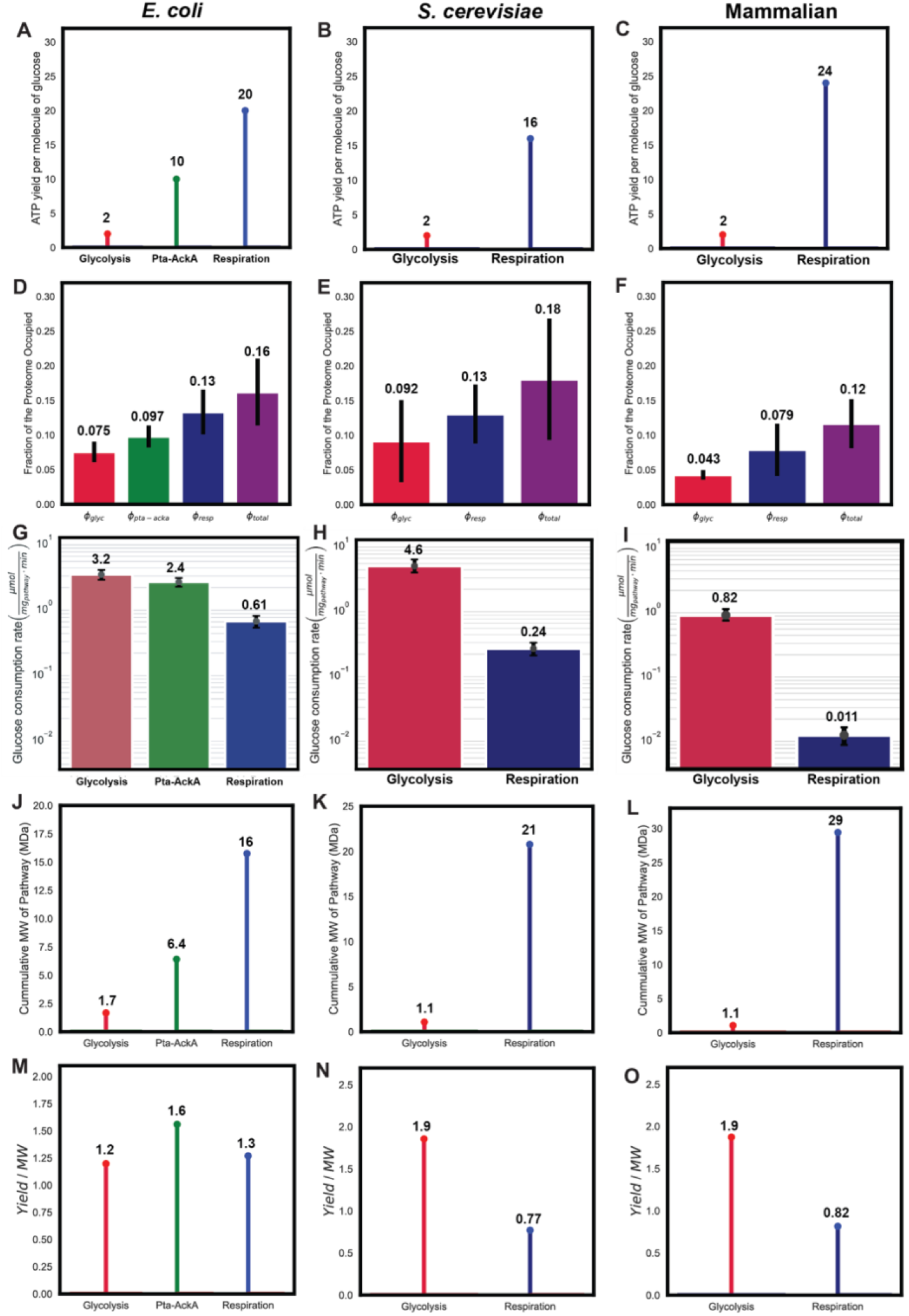
Parameter estimates for *E. coli*, *S. cerevisiae*, and mammalian cells. **A,B,C)** ATP yield per molecule of glucose for glycolysis (red) and respiration (blue) for *E. coli*, *S. cerevisiae*, and mammalian cells, respectively. **D,E,F)** The fraction of the proteome occupied by enzymes for glycolysis (red), respiration (blue), and both pathways (purple) for *E. coli*, *S. cerevisiae*, and mammalian cells, respectively. Error bars are standard deviation. **G,H,I)** Maximal observed glucose uptake rate (μmol per mg cellular protein per min) for glycolysis (red) and respiration (blue) for *E. coli*, *S. cerevisiae*, and mammalian cells, respectively. Error bars are 95 percent confidence interval calculated from 10000 bootstrap iterations. **J, K, L)** Cumulative molecular weight (MDa) of each Pathway **M,N,O)** Ratio of the ATP yield per molecule of glucose to molecular weight (MDa) for each pathway.

**Fig. S5.**
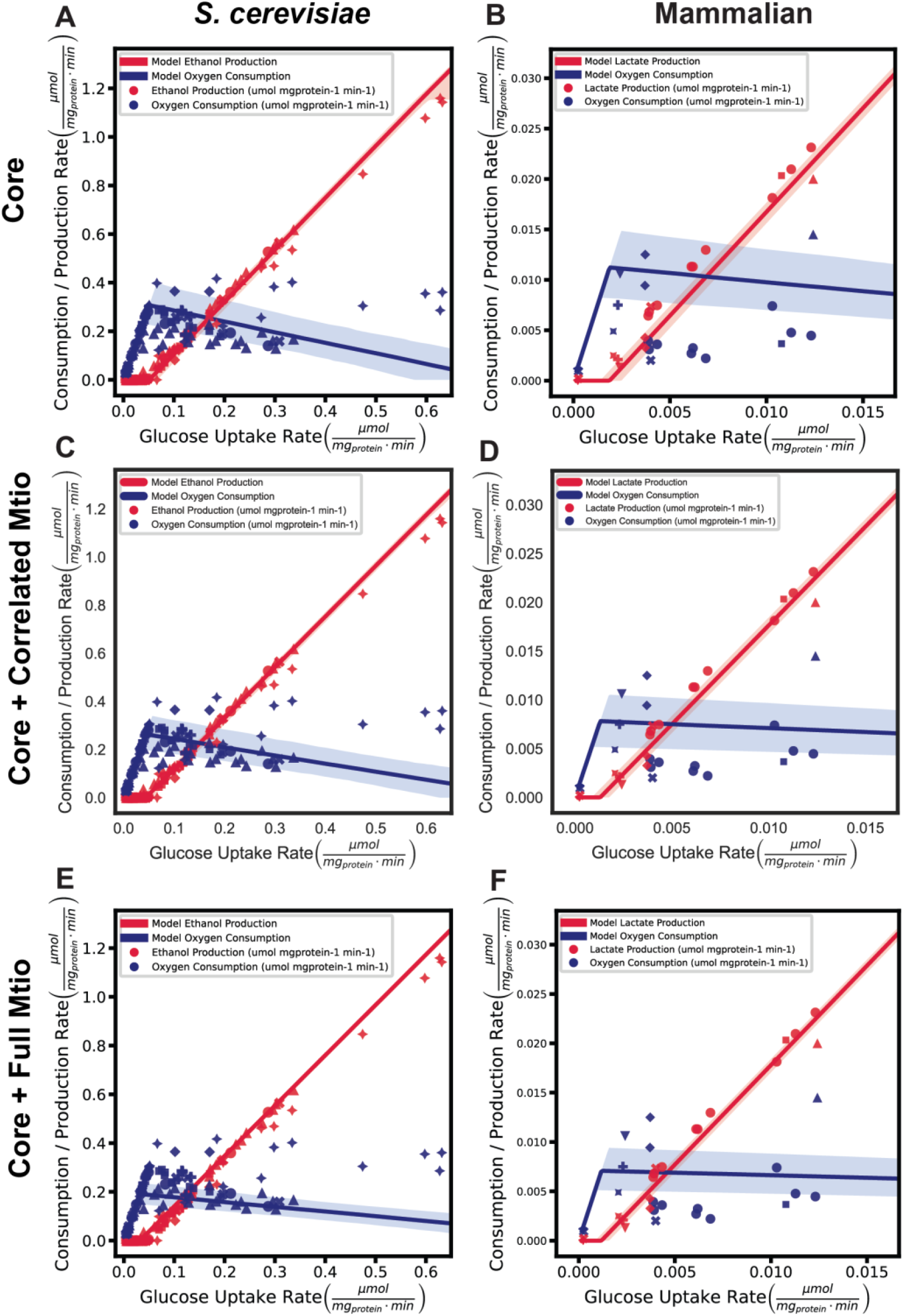
Organism-specific model predictions of ATP Production. **A-F)** Model prediction of glycolysis and respiration rates without any additional mitochondrial proteins (A,B), with inclusion of positively and significantly correlated mitochondrial proteins from Figure 3D,E in the main text (C,D), and all mitochondrial proteins (E,F) for *S. cerevisiae* (A,C,E) and mammalian cells (B,D,F).

## References and Notes

1. O. Warburg, F. Wind, E. Negelein, The metabolism of tumors in the body. J Gen Physiol. 8, 519–530 (1927).

2. O. Warburg, On the origin of cancer cells. Science. 123, 309–314 (1956).

3. M. Potter, E. Newport, K. J. Morten, The Warburg effect: 80 years on. Biochem Soc Trans. 44, 1499–1505 (2016).

4. M. V. Liberti, J. W. Locasale, The Warburg Effect: How Does it Benefit Cancer Cells? Trends Biochem Sci. 41, 211–218 (2016).

5. R. J. DeBerardinis, N. S. Chandel, We need to talk about the Warburg effect. Nature Metabolism. 2, 127–129 (2020).

6. P. Vaupel, G. Multhoff, Revisiting the Warburg effect: historical dogma versus current understanding. The Journal of Physiology. 599, 1745–1757 (2021).

7. J. Fan, J. J. Kamphorst, R. Mathew, M. K. Chung, E. White, T. Shlomi, J. D. Rabinowitz, Glutamine-driven oxidative phosphorylation is a major ATP source in transformed mammalian cells in both normoxia and hypoxia. Molecular Systems Biology. 9, 712 (2013).

8. C.-H. Yao, R. Wang, Y. Wang, C.-P. Kung, J. D. Weber, G. J. Patti, Mitochondrial fusion supports increased oxidative phosphorylation during cell proliferation. eLife. 8, e41351 (2019).

9. K. Brand, Aerobic Glycolysis by Proliferating Cells: Protection against Oxidative Stress at the Expense of Energy Yield, 10.

10. K. A. Brand, U. Hermfisse, Aerobic glycolysis by proliferating cells: a protective strategy against reactive oxygen species1. The FASEB Journal. 11, 388–395 (1997).

11. E. A. Newsholme, B. Crabtree, M. S. Ardawi, The role of high rates of glycolysis and glutamine utilization in rapidly dividing cells. Biosci. Rep. 5, 393–400 (1985).

12. M. G. Vander Heiden, L. C. Cantley, C. B. Thompson, Understanding the Warburg Effect: The Metabolic Requirements of Cell Proliferation. Science. 324, 1029–1033 (2009).

13. Z. Dai, A. A. Shestov, L. Lai, J. W. Locasale, A Flux Balance of Glucose Metabolism Clarifies the Requirements of the Warburg Effect. Biophysical Journal. 111, 1088–1100 (2016).

14. A. Luengo, Z. Li, D. Y. Gui, L. B. Sullivan, M. Zagorulya, B. T. Do, R. Ferreira, A. Naamati, A. Ali, C. A. Lewis, C. J. Thomas, S. Spranger, N. J. Matheson, M. G. Vander Heiden, Increased demand for NAD+ relative to ATP drives aerobic glycolysis. Molecular Cell. 81, 691–707.e6 (2021).

15. Y. Wang, E. Stancliffe, R. Fowle-Grider, R. Wang, C. Wang, M. Schwaiger-Haber, L. P. Shriver, G. J. Patti, Saturation of the mitochondrial NADH shuttles drives aerobic glycolysis in proliferating cells. Molecular Cell. 82, 3270–3283.e9 (2022).

16. E. Rozpędowska, L. Hellborg, O. P. Ishchuk, F. Orhan, S. Galafassi, A. Merico, M. Woolfit, C. Compagno, J. Piškur, Parallel evolution of the make–accumulate–consume strategy in Saccharomyces and Dekkera yeasts. Nat Commun. 2, 302 (2011).

17. T. Pfeiffer, S. Schuster, S. Bonhoeffer, Cooperation and Competition in the Evolution of ATP-Producing Pathways. Science. 292, 504–507 (2001).

18. S. Schuster, T. Pfeiffer, D. A. Fell, Is maximization of molar yield in metabolic networks favoured by evolution? Journal of Theoretical Biology. 252, 497–504 (2008).

19. M. Szenk, K. A. Dill, A. M. R. de Graff, Why Do Fast-Growing Bacteria Enter Overflow Metabolism? Testing the Membrane Real Estate Hypothesis. Cell Systems. 5, 95–104 (2017).

20. D. Molenaar, R. van Berlo, D. de Ridder, B. Teusink, Shifts in growth strategies reflect tradeoffs in cellular economics. Molecular Systems Biology. 5, 323 (2009).

21. M. Basan, S. Hui, H. Okano, Z. Zhang, Y. Shen, J. R. Williamson, T. Hwa, Overflow metabolism in E. coli results from efficient proteome allocation. Nature. 528, 99–104 (2015).

22. A. Flamholz, E. Noor, A. Bar-Even, W. Liebermeister, R. Milo, Glycolytic strategy as a tradeoff between energy yield and protein cost. Proceedings of the National Academy of Sciences. 110, 10039–10044 (2013).

23. J. M. S. Lemons, X.-J. Feng, B. D. Bennett, A. Legesse-Miller, E. L. Johnson, I. Raitman, E. A. Pollina, H. A. Rabitz, J. D. Rabinowitz, H. A. Coller, Quiescent Fibroblasts Exhibit High Metabolic Activity. PLoS Biol. 8, e1000514 (2010).

24. W. Dong, M. A. Keibler, S. J. Moon, P. Cho, N. Liu, C. J. Berrios, J. K. Kelleher, H. D. Sikes, O. Iliopoulos, J. L. Coloff, M. G. V. Heiden, G. Stephanopoulos, Oncogenic metabolic rewiring independent of proliferative control in human mammary epithelial cells (2022), p. 2022.04.08.486845,, doi:10.1101/2022.04.08.486845.

25. M. T. Crow, M. J. Kushmerick, Chemical energetics of slow- and fast-twitch muscles of the mouse. Journal of General Physiology. 79, 147–166 (1982).

26. Y. Chinchore, T. Begaj, D. Wu, E. Drokhlyansky, C. L. Cepko, Glycolytic reliance promotes anabolism in photoreceptors. eLife. 6, e25946 (2017).

27. P. P. Dennis, H. Bremer, Macromolecular composition during steady-state growth of Escherichia coli B-r. J Bacteriol. 119, 270–281 (1974).

28. E. Martínez-Salas, J. A. Martín, M. Vicente, Relationship of Escherichia coli density to growth rate and cell age. Journal of Bacteriology. 147, 97–100 (1981).

29. A. G. Marr, Growth rate of Escherichia coli. MICROBIOL. REV. 55, 18 (1991).

30. E. Bosdriesz, D. Molenaar, B. Teusink, F. J. Bruggeman, How fast-growing bacteria robustly tune their ribosome concentration to approximate growth-rate maximization. The FEBS Journal. 282, 2029–2044 (2015).

31. N. Ertugay, H. Hamamci, Continuous cultivation of bakers’ yeast: Change in cell composition at different dilution rates and effect of heat stress on trehalose level. Folia Microbiol. 42, 463–467 (1997).

32. D. P. Clark, The fermentation pathways of Escherichia coli. FEMS Microbiology Letters. 63, 223–234 (1989).

33. C. R. Dittrich, R. V. Vadali, G. N. Bennett, K.-Y. San, Redistribution of Metabolic Fluxes in the Central Aerobic Metabolic Pathway of E. coli Mutant Strains with Deletion of the ackA-pta and poxB Pathways for the Synthesis of Isoamyl Acetate. Biotechnology Progress. 21, 627–631 (2005).

34. B. Enjalbert, P. Millard, M. Dinclaux, J.-C. Portais, F. Létisse, Acetate fluxes in Escherichia coli are determined by the thermodynamic control of the Pta-AckA pathway. Sci Rep. 7, 42135 (2017).

35. A. S. Divakaruni, M. D. Brand, The Regulation and Physiology of Mitochondrial Proton Leak. Physiology. 26, 192–205 (2011).

36. D. W. Erickson, S. J. Schink, V. Patsalo, J. R. Williamson, U. Gerland, T. Hwa, A global resource allocation strategy governs growth transition kinetics of Escherichia coli. Nature. 551, 119–123 (2017).

37. C. You, H. Okano, S. Hui, Z. Zhang, M. Kim, C. W. Gunderson, Y.-P. Wang, P. Lenz, D. Yan, T. Hwa, Coordination of bacterial proteome with metabolism by cyclic AMP signalling. Nature. 500, 301–306 (2013).

## References and Notes

1. K. M. Carroll, D. M. Simpson, C. E. Eyers, C. G. Knight, P. Brownridge, W. B. Dunn, C. L. Winder, K. Lanthaler, P. Pir, N. Malys, D. B. Kell, S. G. Oliver, S. J. Gaskell, R. J. Beynon, Mol Cell Proteomics, in press, doi:10.1074/mcp.M111.007633.

2. A. Finka, P. Goloubinoff, Proteomic data from human cell cultures refine mechanisms of chaperone-mediated protein homeostasis. Cell Stress and Chaperones. 18, 591–605 (2013).

3. J. R. Wiśniewski, M. Y. Hein, J. Cox, M. Mann, A “Proteomic Ruler” for Protein Copy Number and Concentration Estimation without Spike-in Standards *. Molecular & Cellular Proteomics. 13, 3497–3506 (2014).

4. P. C. Hinkle, P/O ratios of mitochondrial oxidative phosphorylation. Biochim Biophys Acta. 1706, 1–11 (2005).

5. S. Steigmiller, P. Turina, P. Gräber, The thermodynamic H+/ATP ratios of the H+- ATPsynthases from chloroplasts and Escherichia coli. Proceedings of the National Academy of Sciences. 105, 3745–3750 (2008).

6. J. Petersen, K. Förster, P. Turina, P. Gräber, Comparison of the H+/ATP ratios of the H+- ATP synthases from yeast and from chloroplast. Proceedings of the National Academy of Sciences. 109, 11150–11155 (2012).

